# HIV-linked gut dysbiosis associates with cytokine production capacity in viral-suppressed people living with HIV

**DOI:** 10.1101/2022.04.21.489050

**Authors:** Yue Zhang, Sergio Andreu-Sánchez, Nadira Vadaq, Daoming Wang, Vasiliki Matzaraki, Wouter van der Heijden, Ranko Gacesa, Rinse K Weersma, Alexandra Zhernakova, Linos Vandekerckhove, Quirijn de Mast, Leo A. B. Joosten, Mihai G. Netea, Andre van der Ven, Jingyuan Fu

## Abstract

People living with HIV (PLHIV) are exposed to chronic immune dysregulation, even when virus replication is suppressed by antiretroviral therapy (ART). Given the emerging role of the gut microbiome in immunity, we hypothesized that the gut microbiome may be related to the cytokine production capacity of PLHIV. To test this hypothesis, we collected metagenomic data from 143 ART-treated PLHIV and assessed the *ex vivo* production capacity of eight different cytokines (IL-1β, IL-6, IL-1Ra, IL-10, IL17, IL22, TNF and IFN-γ) in response to different stimuli. We also characterized CD4^+^ T cell–counts, HIV reservoir and other clinical parameters. Compared to 190 age- and sex-matched controls and a second independent control cohort, PLHIV showed microbial dysbiosis that was correlated with viral reservoir levels, cytokine production capacity and sexual behavior. Notably, we identified two genetically different *P. copri* strains that were enriched in either PLHIV or healthy controls. The control-enriched strain was negatively associated with IL-10, IL-6 and TNF production, independent of age, sex and sexual behavior, and positively associated with CD4^+^ T cell–level, whereas the PLHIV-enriched strain showed no associations. Our findings suggest that modulating the gut microbiome may be a strategy to modulate immune response in PLHIV.

**Novel Points:**

1. We identified compositional and functional changes in the gut microbiome of PLHIV that were strongly related to sexual behavior.
2. HIV-associated bacterial changes are negatively associated with HIV reservoir. The relative abundance of *Firmicutes bacterium CAG 95* and *Prevotella sp CAG 5226* both show a negative association with CD4^+^ T cell–associated HIV-1 DNA.
3. *Prevotella copri* and *Bacteroides vulgatus* show association with PBMC production capacity of IL-1β and IL-10 that is independent of age, sex, BMI and sexual behavior.
4. We observed two genetically different *P. copri* strains that are enriched in PLHIV and healthy individuals, respectively.
5. The control-related *P. copri* strain specifically shows a negative association with IL-10, IL-6 and TNF production and a positive association with CD4^+^ T cell–level. This suggests it plays a potential protective role in chronic inflammation, which may be related to enrichment of a specific epitope peptide.

## Introduction

Human Immunodeficiency Virus (HIV) infection induces chronic activation of the innate and adaptive immune systems, leading to a chronic inflammatory state ^1^. Combination antiretroviral treatment (ART) significantly decreases immune activation and systemic inflammation but does not restore the homeostasis in the immune system to that seen in healthy populations ^2^. The persistent inflammation, due in part to a dysbalanced cytokine network, contributes to a higher risk of non-AIDS-related morbidity in PLHIV, including cardiovascular disease, neurocognitive disease and certain HIV-related cancers ^2,3^. Previous studies have shown that HIV triggers the production of proinflammatory cytokines (tumor necrosis factor (TNF), interleukin (IL)-6, and IL-1) and anti-inflammatory cytokines (IL-10) ^4,5^. Starting ART has been shown to decrease plasma concentrations of IL-10 and IL-6, but the concentrations of TNF and other proinflammatory cytokines remain elevated ^4^. There are multiple potential causes for this dysregulated cytokine system, such as the effect of HIV, lymphoid tissue damage ^6^ and, in particular, gut dysbiosis ^7^.

HIV infection induces significant changes in gut microbial composition and metabolic function ^8,9^. ART can partially restore the HIV-associated gut dysbiosis, but it cannot normalize the gut microbiome to a pattern resembling that of a healthy control population ^10,11^. Compared with healthy controls, long-term treated PLHIV still exhibit decreased alpha diversity, increased abundances of Enterobacteriaceae and decreased abundances of *Bacteroidetes, Alistipes* ^7^ and butyrate-producing bacteria that help maintain healthy gut homeostasis ^12^. However, no consistent pattern of gut dysbiosis has been defined in long-term treated PLHIV ^13^. One major reason for this is that sexual behavior may have a strong influence on gut microbiota, and most previous studies were not able to control for this factor ^13^. For example, an over-representation of *Prevotella* accompanied with a decrease in *Bacteroides* in PLHIV was recently found to be due to men who have sex with men (MSM) status rather than HIV infection ^14–16^. Considering that a major part of the HIV-infected population in Europe and North America is composed of MSM, the biological importance of this feature needs to be explored.

In long-term treated PLHIV, a link between gut dysbiosis and host cytokine levels has been reported. For instance, in PLHIV with atherosclerosis, class Clostridia was positively correlated with plasma levels of IL-1β and IFN-γ ^17^, while coproic acid, a gut bacteria–derived short-chain fatty acid (SCFA), was linked with decreased expression of IL-32 ^18^. Mechanistically, the bacteria–cytokine association is potentially based on bacterial components and products ^9,19–21^. For example, heat-killed *Escherichia coli* induced higher production of IL-17 and IFN-γ in HIV-exposed mononuclear cells *ex vivo* ^19^. In addition, lipopolysaccharide (LPS), a Gram-negative bacterial cell wall component, induced a depletion of CD4^+^ cells by increasing expression of HIV coreceptor C-C chemokine receptor type 5 on CD4^+^ cells ^20^. In addition, butyrate, a product of saccharolytic fermentation of dietary fibers by gut microbiota, decreased gut T cell activation in an *ex vivo* human intestinal cell culture model ^21^. The downregulation of anti-inflammatory bacterial pathways, such as SCFA biosynthesis or indole production, also contributes to gut inflammation in long-term treated PLHIV ^9^.

The present study documents a detailed profile of gut microbial composition and function at both species- and strain-level using metagenomic sequencing. We then characterize the role of gut dysbiosis in relation to HIV clinical phenotypes and PBMC cytokine production capacity. Notably, the association between cytokine production capacity and gut microbiome has only been studied in healthy populations ^22^ and in a limited number of PLHIV ^10^. In addition, we control for sexual behavior–related factors in the association analysis. Finally, we identify two *P. copri* strains with different genetic repertoires that exhibit enrichment in PLHIV and HCs, respectively. One of these two *P. copri* strains showed associations with the PBMC production capacity of IL-10, IL-6 and TNF, as well as CD4^+^ T cell–level.

## Results

### Microbial dysbiosis in PLHIV

#### Differences in microbial composition and function

The present study included 143 PLHIV and 190 healthy individuals with matched age and sex from the Dutch Microbiome Project (DMP) cohort (hereafter referred to as matched healthy controls (HCs)). Participants’ baseline characteristics are shown in Supplementary Table 1. Included PLHIV were on long-term ART (median 6.35 years) and were virally suppressed (plasma HIV-RNA < 200 copies/mL). However, when compared to the matched HCs, the gut microbial composition of the PLHIV still showed a significant decrease in alpha diversity (species-level Shannon index, Wilcoxon rank-sum test, P = 2.4×10^−5^, Fig. 1a) and a distinct composition that was reflected by a significant difference in beta diversity (PERMANOVA, P < 1.0×10^−3^, R^2^ = 0.07, Fig. 1b), even when adjusting for body mass index (BMI) and read counts. These observations are consistent with findings of previous studies ^23–25^. With the aid of metagenomics data, we also observed that PLHIV showed a lower diversity of bacterial metabolic pathways (pathway-level Shannon index, Wilcoxon rank-sum test P = 4.0×10^−4^, Supplementary Fig. 1) and different functional profiles (PERMANOVA, P < 1.0×10^−3^, R^2^ = 0.04, Fig. 1c). To explore the bacterial taxa and pathways that showed significantly different abundance in PLHIV, we confined the differential abundance analysis to the 123 common species and 331 metabolic pathways present in ≥ 20% of samples in at least one cohort. A linear regression model with correction for read counts, BMI and smoking status revealed 76 (62.6%) species and 163 pathways (49.2%) with differential abundances between PLHIV and HCs (False Discovery Rate (FDR) < 0.05, Supplementary Tables 2–3).

**Fig. 1.**
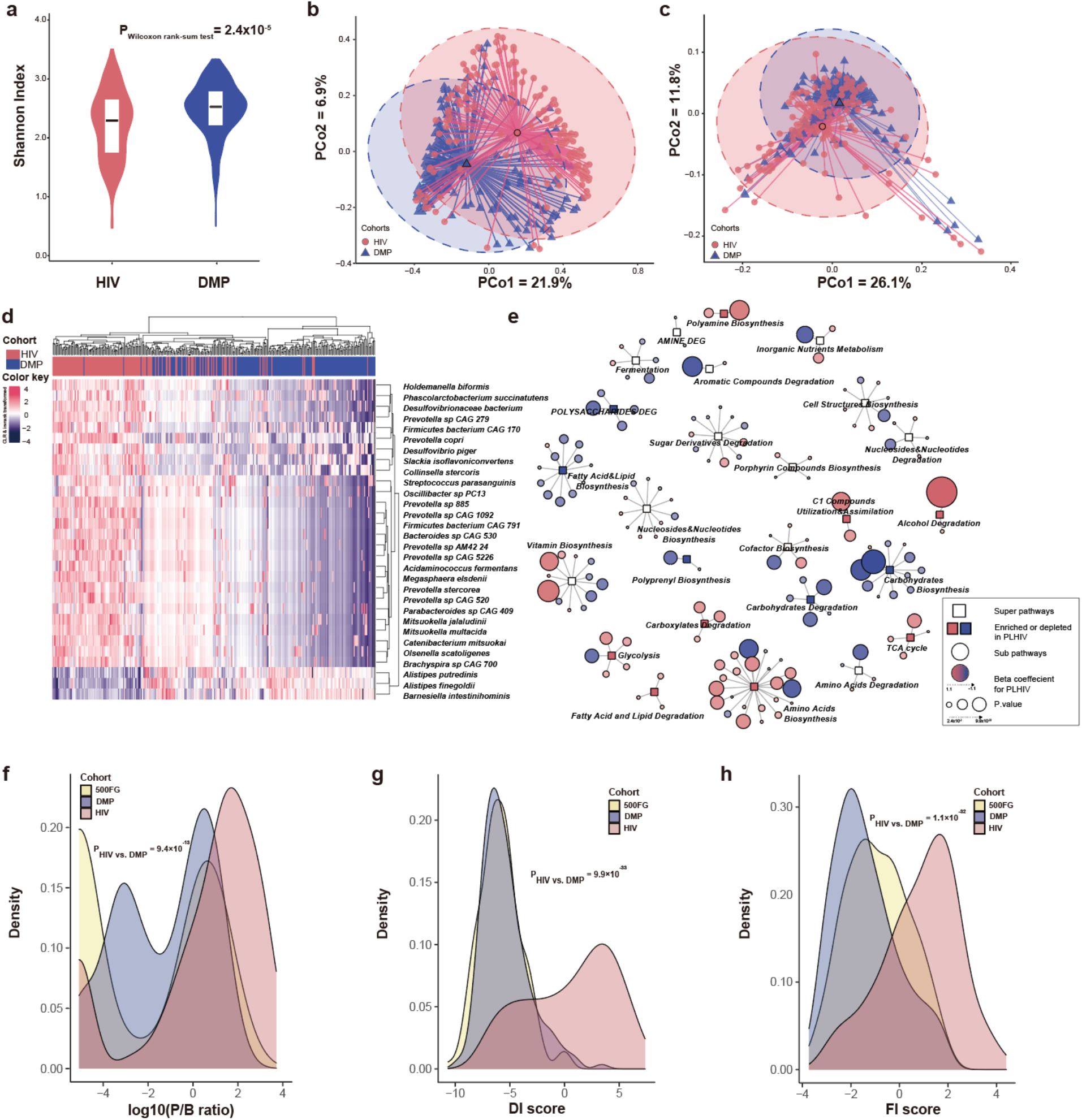
PLHIV show a distinct gut microbiome composition and function compared to HCs. **a**. Comparison of microbial alpha diversity between the PLHIV cohort and HCs from the DMP cohort. Y-axis refers to the Shannon index at the species level. **b–c**. Beta diversity based on Bray-Curtis distance of species and pathway abundance is shown in a principal coordinates analysis (PCoA) plot with centroids for PLHIV and HCs. The coordinates of the centroids are set as the mean value of the principal components for each cohort. **d**. Heatmap depicting the relative abundance of the top 30 species that differed significantly between PLHIV and HCs. Data is centered log-ratio (CLR)-transformed and then inverse-rank transformed to follow a normal distribution. Differentially abundant species were selected using a linear regression model with correction for BMI, read counts and smoking status. **e**. Network of bacterial pathways that were significantly different between PLHIV and HCs. Rectangular nodes represent super-pathways, with the colors showing enrichment (pink) or depletion (blue) in PLHIV. Circular nodes show sub-pathways belonging to the super-pathways. The color of circles shows the log2 value of fold change between the relative abundance of pathway in PLHIV and HCs, where a gradient is applied depending on foldchange. Circle size indicates p-value. Lines connect each pathway to its respective super-pathway. Only super-pathways including two or more pathways and sub-pathways with FDR < 0.05 are shown. Differentially abundant pathways were selected using a linear regression model with correction for BMI, read counts and smoking status. **f–h**. Density curves of the P/B ratio, DI score and FI score for the three cohorts depicting the different distribution of these bacterial signatures in these cohorts. Significance was tested using Dunn’s test.

#### Differentially abundant species

57 of the 76 differentially abundant species showed enrichment in PLHIV. Increased abundances of *Prevotella* and *Prevotellaceae* in PLHIV were widely observed by previous studies using 16S rRNA sequencing ^8^. Our metagenomics-based analysis further identified eight *Prevotella* species enriched in PLHIV (Supplementary Table 2). The most significant was *Prevotella sp 885*, which showed a 3.8-fold increase in relative abundance (Linear regression, P = 2.1×10^−27^), followed by *Prevotella sp CAG 520* with an 8.0-fold increase (P = 5.4×10^−24^) and *Prevotella sp CAG 1092* with a 4.2-fold increase (P = 3.4×10^−23^, Fig. 1d, Supplementary Fig. 2). Also consistent with other studies ^10,26^, we found an increase of *Desulfovibrionaceae bacterium* (P = 7.8×10^−24^) and *Megasphaera elsdenii* (P = 3.9×10^−23^). *Megasphaera* species, as members of the vaginal microbiome, were associated with a higher risk of acquiring HIV in a prospective study of HIV-infected South African women ^27^. The abundances of 19 species were decreased in PLHIV, including species from the previously reported species *Bacteroides* and *Alistipes* ^7,8^: *B. ovatus, B. uniformis, B. vulgatus, A. finegoldii* and *A. putredinis*. We also identified several novel HIV-associated species, including *Barnesiella intestinihominis*, which was mostly depleted in PLHIV (P = 7.5×10^−15^). This bacterium was previously identified as an “oncomicrobiotic” due to its capacity to promote the infiltration of IFN-γ-producing γδT cells in cancer lesions, which can ameliorate the efficacy of the anti-cancer immunomodulatory agent cyclophosphamide ^28^.

#### Differentially abundant pathways

At metabolic pathway–level, we observed differential abundances in several amino acid biosynthesis pathways, including enriched L-tryptophan biosynthesis (PWY-6629) and depleted L-ornithine and L-citrulline biosynthesis pathways (ARGININE-SYN4-PWY, CITRULBIO-PWY) in PLHIV (Supplementary Table 3). Importantly, tryptophan and citrulline both play critical roles in inflammation ^29,30^, while ornithine can later be turned into Nitric Oxide (NO), which is important for vascular function ^31,32^. In addition, the reductive tricarboxylic acid (TCA) cycle I (P23-PWY) was enriched in PLHIV. The reductive TCA cycle is a carbon dioxide–fixation pathway significant for the production of organic molecules for the biosynthesis of sugars, lipids, amino acids and pyrimidines ^33^. Moreover, PLHIV showed lower abundances of bacterial pathways involved in fatty acid, lipid and carbohydrate biosynthesis (Fig. 1e), suggesting a reduced capacity of the microbiome to digest certain nutritional elements.

#### Dysbiosis index

Altogether, our data show dysbiosis in both gut microbial composition and metabolic function in PLHIV. Previous studies have suggested the *Prevotella*-to-*Bacteroides* ratio (P/B ratio) as a landmark parameter for PLHIV ^34,35^, and our finding of a significantly higher P/B ratio in PLHIV compared to HCs confirms these observations (Dunn’s test, P = 9.4×10^−13^, Fig. 1f, Supplementary Table 4, Supplementary Fig. 3a). We also constructed a Dysbiosis Index (DI) based on the differentially abundant species by calculating the log2-ratio of the geometric mean of PLHIV-enriched species (57 species) to PLHIV-depleted species (19 species). This DI score was significantly higher in PLHIV than in HCs (P = 9.9×10^−33^, Fig. 1g, Supplementary Table 4). We then sought to validate the DI score in an independent cohort with similar metagenomic data and the same DNA isolation method. While no such data were available for an independent cohort of PLHIV, data was available for a separate healthy cohort: 500FG (500 Functional Genomics) (Supplementary Table 1). We found that the DI score of 500FG was not different from the DMP controls (P = 0.29, Supplementary Fig. 3b), but was significantly lower than that of our cohort of PLHIV (P = 9.6×10^−37^). We also constructed a Function Imbalance (FI) score using the log2-ratio of the geometric mean of PLHIV-enriched bacterial pathways (87 pathways) to PLHIV-depleted bacterial pathways (76 pathways) (Fig. 1h). This FI score was also significantly higher in PLHIV than in DMP HCs (P = 1.1×10^−32^, Supplementary Table 4, Supplementary >Fig. 3c), and this was supported by the observation that the 500FG FI was also significantly lower than that of PLHIV (P = 6.7×10^−18^).

### HIV-associated gut dysbiosis associates with clinical phenotypes

We conducted a systematic association between HIV-related parameters and microbial alpha diversity, beta diversity, P/B ratio, DI score and FI score, correcting for sex, age and read counts (Supplementary Fig. 4 and 5, Supplementary Table 5). HIV clinical parameters were also taken into account, including the time between HIV diagnosis and inclusion in the study or cART initiation, CD4^+^ T cell counts (nadir and latest, recovery after cART), plasma viral loads and HIV-1 reservoir measurements in circulating CD4^+^ T cells, including CD4^+^ T cell–associated HIV-1 DNA (CA-HIV-DNA) and CD4^+^ T cell–associated HIV-1 RNA (CA-HIV-RNA) levels. Sexual behavior was also considered part of the HIV clinical phenotype, including the number of sexual partners in the previous year (Num-P) and sexual orientation (SO), e.g. MSM and men who have sex with women, including receptive anal intercourse (RAI).

We did not observe any significant association between the Shannon index of bacterial species and any HIV clinical phenotypes (Supplementary Table 6). However, six parameters were associated with bacterial beta diversity, four with P/B ratio, three with DI score and four with FI score at FDR < 0.1 level (Fig. 2a, Supplementary Table 6). Notably, sexual behavior was among the strongest-associated factors. For example, SO was the top factor positively associated with the P/B ratio (Spearman correlation, P = 5.7×10^−12^) and DI score (P = 6.4×10^−12^), followed by Num-P and RAI (Supplementary Table 6). Moreover, MSM and RAI, as well as larger Num-P, were associated with increased P/B ratio, DI and FI score. We also observed associations for other HIV clinical phenotypes, such as an association between beta diversity and the HIV reservoir parameters CA-HIV-DNA and CA-HIV-RNA levels (P = 6.0×10^−3^ and 8.0×10^−3^, respectively) and an association between P/B ratio and CD4 recovery relative rate (P = 0.02). However, these associations were largely dependent on sexual behavior, as none remained significant after correcting for SO and Num-P (Supplementary Table 6, Supplementary Fig. 6). We further assessed whether HIV clinical parameters were associated with individual species and metabolic pathways. After correcting for age, sex, read counts and sexual behavior, no significant associations were detected for species or pathways, although we did observe suggestive associations for *Firmicutes bacterium CAG 95* and *Prevotella sp CAG 5226* with HIV reservoir measurements at a normal significance level (Fig. 2b–d, Supplementary Table 7). At metabolic pathway level, HIV duration and cART duration tended to be the strongest factors besides sexual behavior factors linked with metabolic pathways (Supplementary Table 8). For example, the pathways of stearate, octanoyl and oleate biosynthesis showed positive correlation with HIV duration, while the top result for cART duration was a negative association with the polyamine biosynthesis pathway (Supplementary Table 8).

**Fig. 2.**
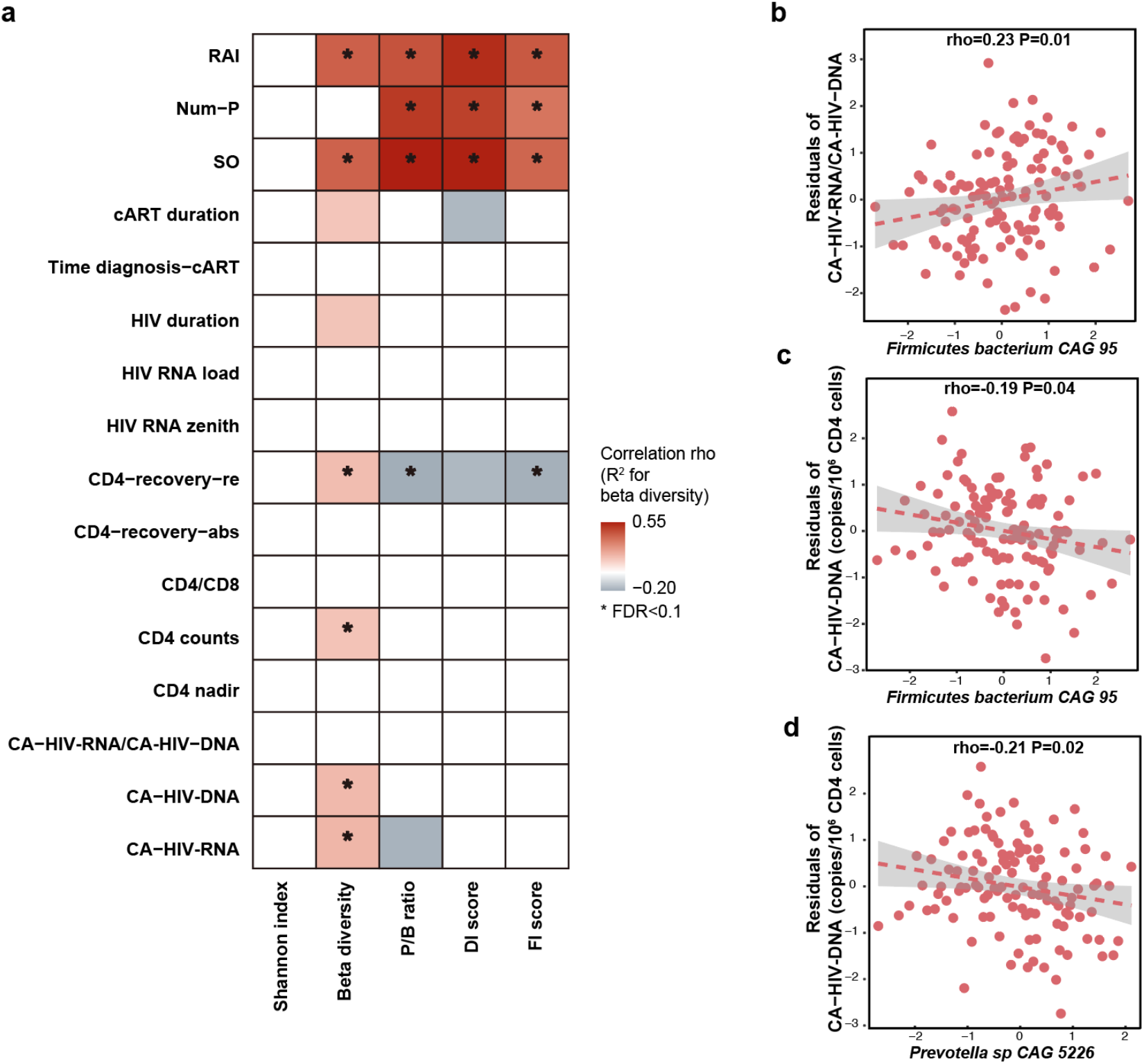
Association between HIV-associated gut dysbiosis and HIV-related phenotypes. **a**. Heatmap depicting the associations between HIV-associated bacterial signature (Shannon index, beta diversity, P/B ratio, DI and FI score) and HIV-related phenotypes, using the Spearman correlation test, with HIV-related phenotypes corrected for age, sex and read counts. Box color indicates Spearman correlation rho. A white box indicates P > 0.05. R^2^ is calculated using PERMANOVA based on Bray-Curtis distance of species and then multiplied by ten to rescale. **b–d**. Associations between individual species and HIV reservoir level, with the HIV reservoir level corrected for age, sex and read counts and sexual behavior.

### *Prevotella copri* and *Bacteroides vulgatus* associate with cytokine production capacity

Despite long-term cART, the immune responses of PLHIV are known to be different than those of healthy controls ^36^. In a previous study from our group ^1^, PBMCs of PLHIV and healthy controls were stimulated *ex vivo* with 12 different microbial stimuli and the cytokine production capacity assessed, including monocyte-derived (IL-1β, IL-6 and TNF), lymphocyte-derived proinflammatory cytokines (IL-17, IL-22 and IFN-γ) and anti-inflammatory cytokines (IL-10 and IL-1 receptor antagonist, IL-1Ra) (Supplementary Table 9). This identified a significant increase in the production of proinflammatory cytokines (e.g. IL-1β and IL-6 and TNF) in PLHIV.^1^ By comparing the cytokine production data of our HIV cohort to 173 samples from the 500FG cohort (different samples from our previous studies ^1,36^), we identified 24 cytokine abundances that were significantly different (Fig. 3a, Supplementary Table 10). PLHIV showed a significant increase of IL-1β, IL-6 and TNF production upon stimulation with Pam3Cys (TLR2 ligand), LPS (TLR4 ligand) and *C. albicans hyphae*, but decreased IFN-γ production upon stimulation with *S. aureus* and *C. albicans hyphae*. Differing cytokine production capacities were also related to some HIV clinical phenotypes. MSM status was associated with increased proinflammatory cytokine responses (e.g. Pam3Cys-induced TNF and *S. aureus*-induced IL-22 production), whereas higher Num-P was linked with decreased anti-inflammatory cytokine responses (e.g. Pam3Cys and LPS-induced IL-10 production, Supplementary Fig. 7, Supplementary Table 11).

**Fig. 3.**
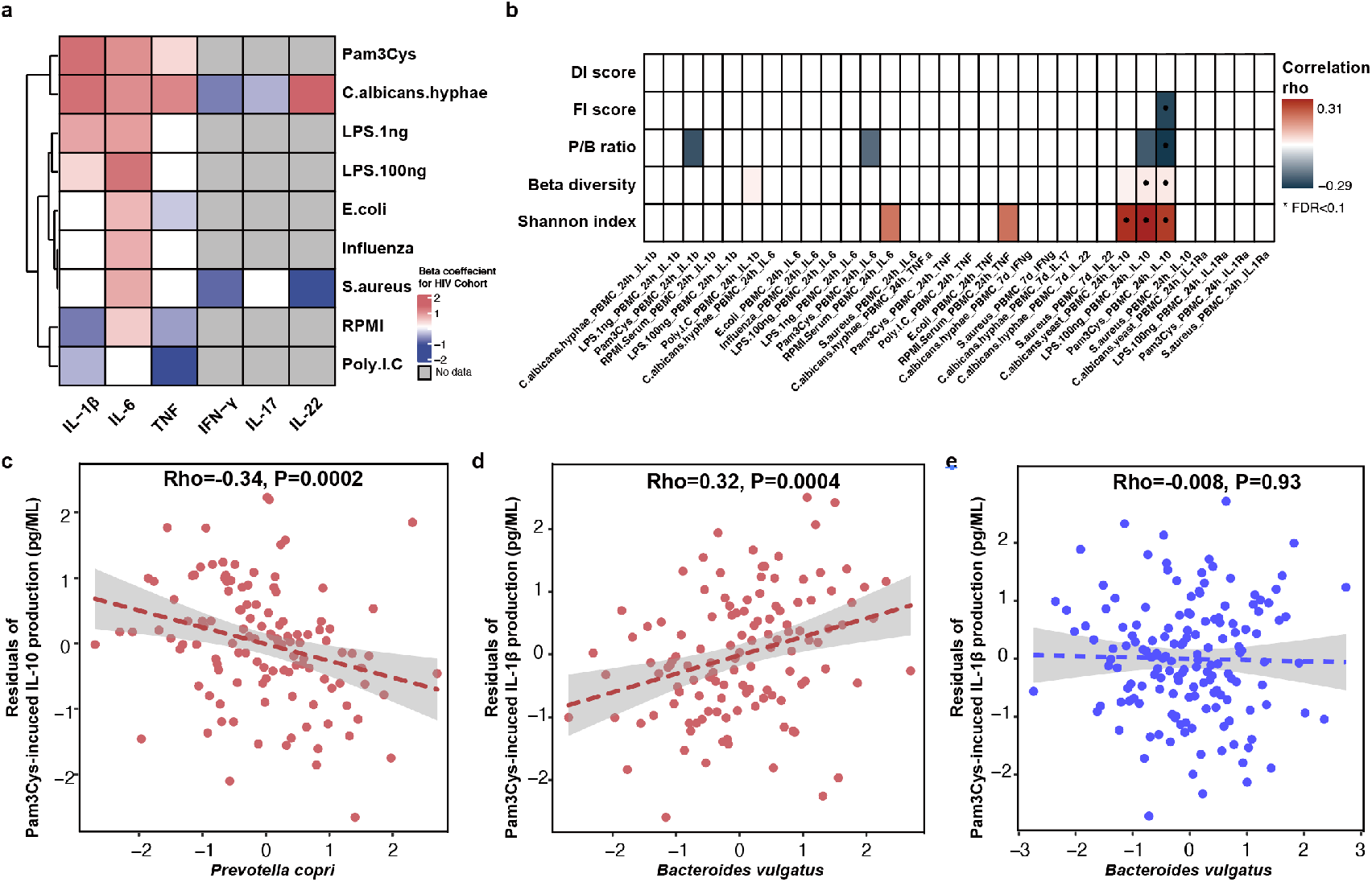
Association between gut microbiome and inflammatory cytokine production. **a**. Heatmap showing *ex vivo* cytokine production enriched (red) and depleted (blue) in PLHIV as compared with HCs from 500FG. A linear regression model (age and sex included as covariates) was used to calculate the P values. Box color indicates the Spearman correlation rho. White box indicates P > 0.05. Gray box indicates no measurement. **b**. Heatmap showing Spearman correlation rho between cytokine production and HIV-associated bacterial signature (Shannon index, beta diversity, P/B ratio, DI and FI scores), with cytokine production corrected for age, sex, read counts and sexual behavior. White box indicates P > 0.05. **c–d**. Association between relative abundance of species and cytokine production in PLHIV: (**c**) *Prevotella copri* with Pam3Cys-induced IL-10 production and (**d**) *Bacteroides vulgatus* with Pam3Cys-induced IL-1β production. The relative abundance of the species is CLR- and inverse-rank transformed. Cytokine production is corrected for age, sex, read counts and sexual behavior. **e**. Association between relative abundance of *Bacteroides vulgatus* and Pam3Cys-induced IL-1β production in HCs from 500FG.

Interestingly, we observed significant associations of cytokine production capacity with microbial alpha and beta diversity and P/B ratio, as well as four associations with DI, FI score and individual species (Supplementary Tables 12 and 13). No associations were observed with metabolic pathway abundance (Supplementary Table 14). In particular, IL-10 production upon stimulation with Pam3Cys or LPS was negatively associated with P/B ratio and FI score but positively associated with bacterial Shannon index (Linear regression, P < 0.05, Fig. 3b, Supplementary Table 12), after correction for age, sex and sexual behavior. At individual species–level, the top association was between Pam3Cys-induced IL-10 production and the relative abundance of *P. copri* (rho = -0.37, P = 9.1×10^−6^, Supplementary Table 13), after correcting for age, sex and read counts. After further adjustment for SO and Num-P, the association between *P. copri* and IL-10 production remained significant (rho = -0.34, P = 1.9×10^−4^, Fig. 3c, Supplementary Table 13). In addition, we also detected a positive significant association between *B. vulgatus* and Pam3Cys-induced IL-1β production (rho = 0.33, P = 7.4×10^−5^), and this remained significant after controlling for age, sex, read counts and sexual behavior (rho = 0.32, P = 3.7×10^−4^, Fig. 3d). Strikingly, this association was not significant in 500FG (rho = -7.8×10^−3^, P = 0.93, Fig. 3e), showing a significant heterogeneity effect (Cochran-Q test, P = 0.003, Supplementary Table 13).

### *Prevotella copri* strains in PLHIV are genetically different

In microbial species, strain-level genomic makeup is critical in determining their functional properties within human bodies ^37^. We therefore wondered whether PLHIV harbor different strains of *P. copri* and *B. vulgatus*, thereby affecting cytokine production capacity. To examine this, we performed an analysis based on the presence or absence of their gene repertoire using PanPhlan3 ^38^. As *B. vulgatus* did not show any distinct clusters according to genetic content, we did not study it further. We generated *P. copri* genetic repertoire profiles for 254 samples from the PLHIV cohort (n = 102), DMP cohort (n = 83) and 500FG cohort (n = 69). Hierarchical clustering analysis based on the Jaccard distance of presence or absence of gene families revealed three distinct clusters at a distance cutoff of 0.4 (Fig. 4a). Cluster 1 was relatively small (n = 13) and was therefore excluded in the following analysis. In contrast, clusters 2 and 3 were larger (n = 137 and 104, respectively). Interestingly, cluster 2 was significantly enriched in healthy individuals and cluster 3 was enriched in PLHIV (Fisher exact test, P = 3.2×10^−25^, Supplementary Table 15), suggesting that the *P. copri* strains are genetically different in PLHIV compared to healthy individuals. We hereafter refer to these *P. copri* strains as the “control-related strain” and “HIV-related strains”. In particular, the HIV-related strain was enriched in MSM of the PLHIV cohort but did not show enrichment in PLHIV with RAI as a sexual behavior (Fisher exact test, P = 0.01 and 0.17, respectively, Supplementary Table 15).

**Fig. 4.**
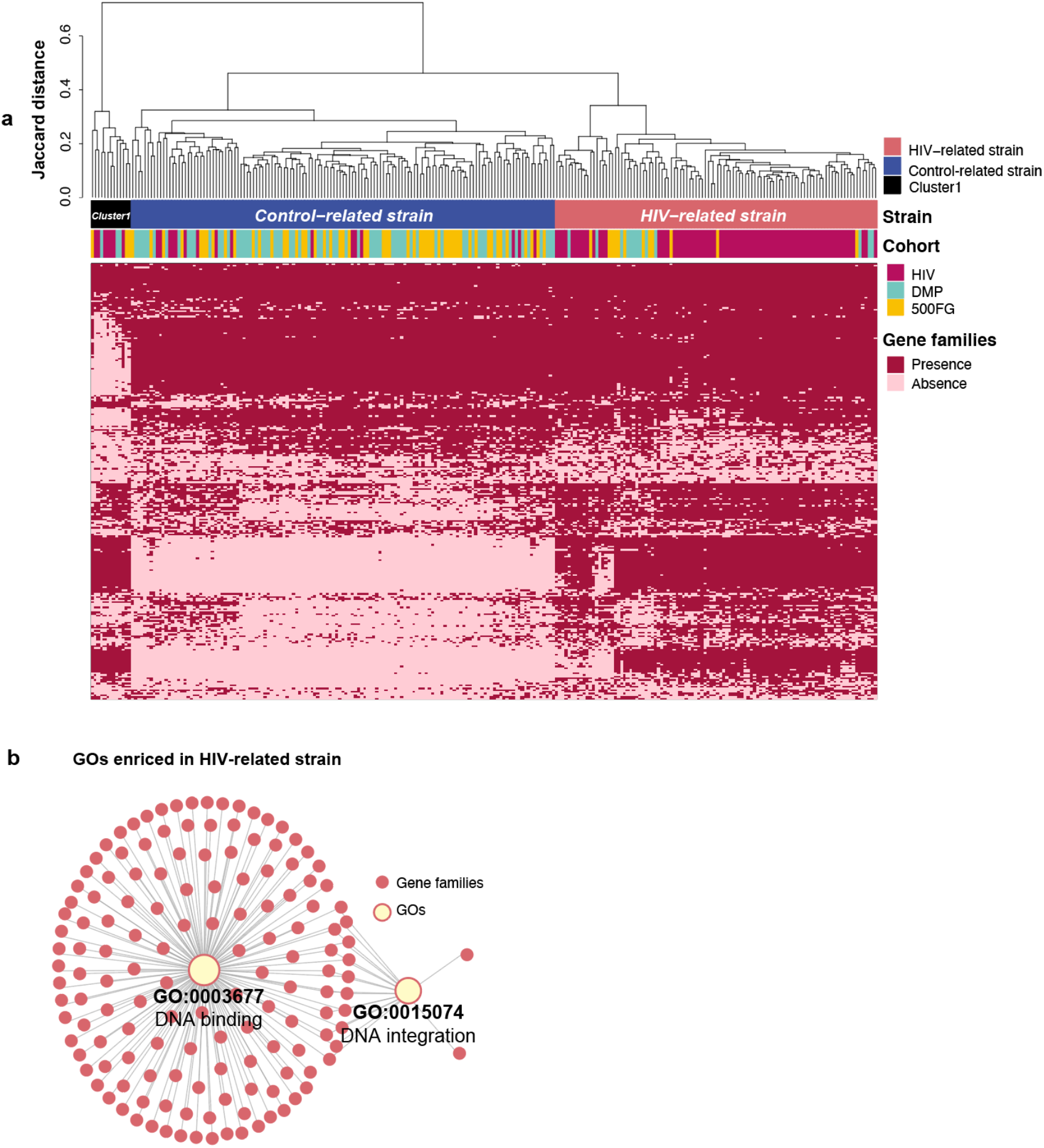
*Prevotella copri* strains with different genetic content. **a.** Heatmap showing the gene family profiles of *P. copri* strains in samples from the three cohorts (254 samples total: 102 PLHIV, 83 DMP and 69 500FG). Each column represents a sample. Each row represents the presence of absence of a gene family. The clustered tree above the heatmap shows the three clusters of *P. copri* strains. Most samples from PLHIV were binned together into the HIV-related strain (right), but 16 samples (middle) showed different profiles and were binned into the HC-related strain. **b.** Network figure showing the two GOs enriched in the HIV-related strain. Pink dots represent different gene families. Yellow dots represent GOs. Lines indicate that the gene families are annotated to the corresponding GO.

When comparing the gene profiles of the two strains, 2,629 out of 4,260 gene families were differentially abundant after correcting for age, sex and read counts (Logistic regression, P < 0.05, FDR < 0.05, Fig. 4a, Supplementary Table 16). We then conducted Gene Ontology (GO) enrichment analysis based on these differential gene families. Two GOs showed enrichment in the HIV-related strain (hypergeometric test, P < 0.05, FDR < 0.05, Fig. 4b, Supplementary Table 17): the molecular function of DNA binding (GO:0003677) and the biological process of DNA integration (GO:0015074). In contrast, no GOs showed enrichment in the control-related strain (Supplementary Table 18).

### *Prevotella copri* strains exhibit distinct immune impacts between PLHIV and HCs

As shown above, the control-related strain and HIV-related strain exhibited distinct genetic content. Likewise, these two strains also showed different associations with cytokine production, and importantly, these associations tended to be influenced by host immune state. The control-related strain showed negative associations with IL-10, IL-6 and TNF production in PLHIV, after correcting for age, sex, read counts and sexual behavior (Linear regression, beta_100ngLPS-induced IL-10_ = -4.7, beta_Pam3Cys-induced IL-6_ =-5.1, beta_100ngLPS-induced IL-6_ = -4.6, beta_No stimulation TNF_ = -3.2; P_100ngLPS-induced IL-10_ = 4.0×10^−4^, P_Pam3Cys-induced IL-6_ = 7.0×10^−3^, P_100ngLPS-induced IL-6_ = 7.2×10^−3^, P_No stimulation TNF_ = 1.7×10^−2^, Fig. 5a–d, Supplementary Table 19). However, in the 500FG cohort, the control-related strain showed less strong and even opposite associations with cytokine production (Linear regression, beta_Pam3Cys-induced IL-6_ = 1.4, beta_100ngLPS-induced IL-6_= 0.69, beta_Nostimulation TNF_ = 0.46; P_Pam3Cys-induced IL-6_ = 3.1×10^−3^, P_100ngLPS-induced IL-6_ = 0.12, P_No stimulation TNF_ = 0.17, Supplementary Fig. 8, Supplementary Table 20). By contrast, no significant association was found for the HIV-related strain in PLHIV and HCs (Supplementary Tables 19 and 20). In summary, we observed a heterogeneity effect in the cytokine association with the control-related and HIV-related strain in PLHIV (Cochran-Q test, Supplementary Table 19), as well as a heterogeneity effect in the cytokine association with the control-related strain between PLHIV and HCs (Supplementary Table 21).

**Fig. 5.**
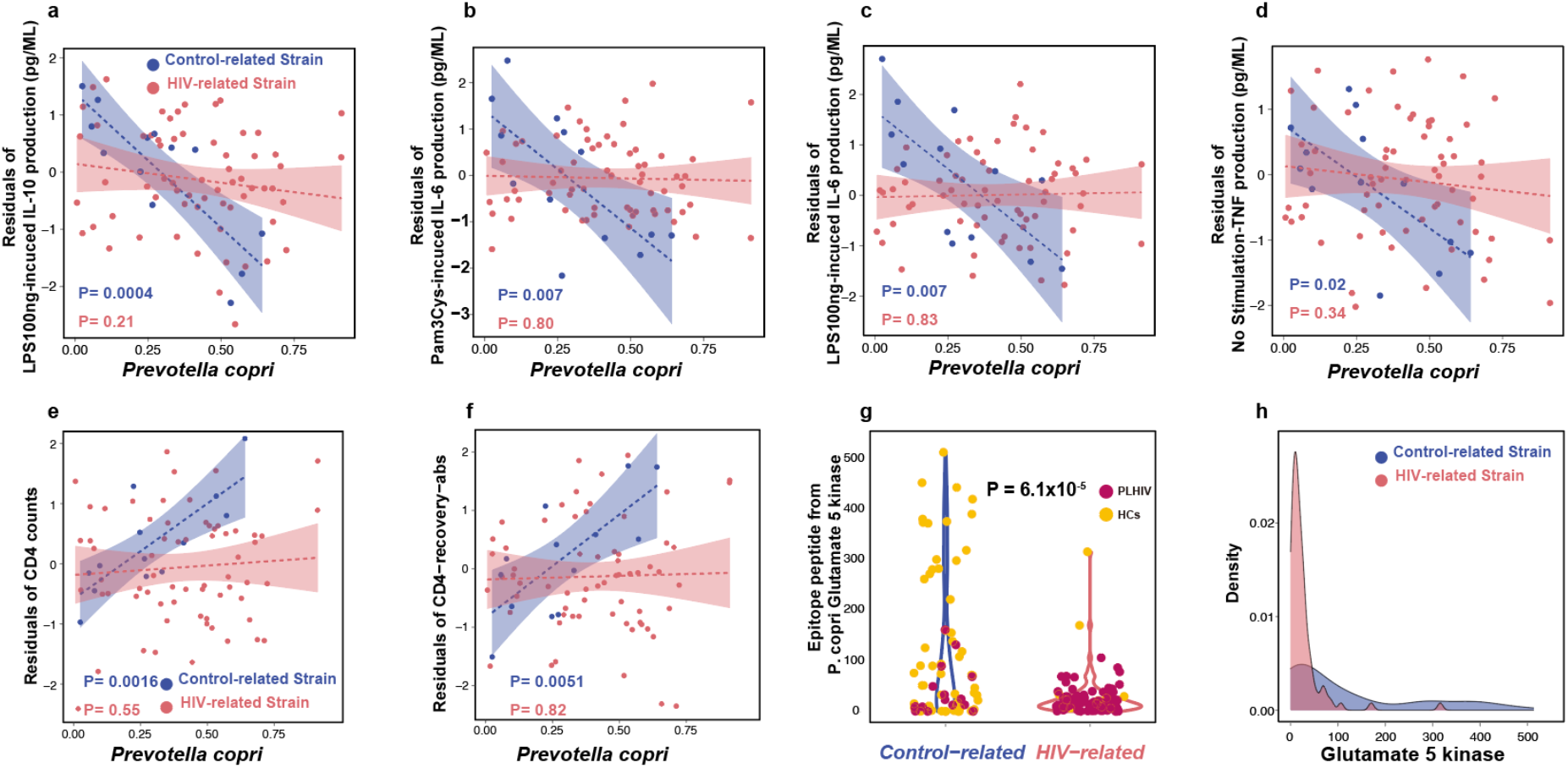
*Prevotella copri* strains with different immune functions. **a–d**. Distinct associations between relative abundance of control-related strain and HIV-related strain with IL-10, IL-6 and TNF production capacity in PLHIV, using linear regression. Cytokine production is corrected for age, sex, read counts and sexual behavior. Blue dots represent the PLHIV with the control-related strain. Dark pink indicates PLHIV with the HIV-related strain. **e–f**. Distinct associations between relative abundance of control-related strain and HIV-related strain with CD4 counts (**e**) and CD4-recovery-abs level (**f**). **g–h**. Violin plot (**g**) and density distribution plot (**h**) of the *P. copri* epitope peptide (from Glutamate 5 kinase) level between the two kinds of strains in PLHIV and HCs from the 500FG cohort. Linear regression was used to test significance, controlling for age, sex and read counts

We also checked the association between *P. copri* strains and HIV-related parameters. The control-related strain showed positive association with CD4^+^ T cell counts (CD4 counts) and CD4^+^ T cell absolute recovery level after cART (CD4-recovery-abs) (beta_CD4 counts_ = 3.2, beta_CD4-recovery-abs_ = 3.5, P_CD4 counts_ = 1.6×10^−3^, P_CD4-recovery-abs_= 5.1×10^−3^, Fig. 5e-f, Supplementary Table 22). These strain–CD4 associations also showed heterogeneity between the control- and HIV-related strains (P = 1.9×10^−3^ and 3.0×10^−3^, respectively, Supplementary Table 22).

To explore the potential mechanism behind the distinct immune functions of the two *P. copri* strains, we searched the Immune Epitope Database and Analysis Resource (IEDB) and found five epitope peptides derived from different *P. copri* proteins (Supplementary Table 23). These peptides can be presented by antigen-presenting cells and induce IFN-γ production by T cells and antibody secretion by B cells. We then compared the abundances of these five peptides between the two *P. copri* strains from PLHIV and 500FG cohort (methods). Interestingly, only one peptide from *P. copri*, glutamate 5-kinase protein, was found to be significantly enriched in the samples with the control-related strain as compared with the samples with the HIV-related strain (Linear regression, P = 6.1×10^−5^, Fig. 5g-h, Supplementary Table 24), suggesting that it is potentially contributing to the immune function of the control-related strain.

## Discussion

We investigated the role of the gut microbiome in functional immune responses during HIV infection using metagenomic sequencing and the cytokine responses of PBMCs upon stimulation, which can profile bacterial composition and functionality and exhibit the functional changes in immunity^1^. Consistent with the literature, we observed a lower alpha diversity and higher P/B ratio in PLHIV, as well as a depletion of *Alistipes* species ^7,8^. In contrast to previous studies, we found some SCFA-producing species and beneficial species to be enriched in PLHIV in our study, including *Acidaminococcus fermentans* ^39^ and *Faecalibacterium prausnitzii* ^40^, as well as the equol-forming species *Slackia isoflavoniconvertens* ^41^ and the anti-tumorigenic species *Holdemanella biformis* ^42^. This inconsistency may be due to the fact that the PLHIV in our cohort had a longer duration of ART intake (median years = 6.4) compared to previous studies, which is supported by earlier evidence that ART can partially restore the gut microbial composition ^10^. Functionally, PLHIV in our study showed increased microbial capacity of L-tryptophan biosynthesis and decreased de novo biosynthesis of ornithine from 2-oxoglutarate, functions that have been related to inflammation and vascular function ^29,32^, respectively. This observation supports the idea that those functional changes in the gut microbiome in PLHIV using ART contribute to the persistent inflammation seen in these individuals.

Infected long-lived memory CD4^+^ T cells make up the majority of cell types constituting the HIV reservoir ^43^. To the best of our knowledge, no other studies have ever reported associations between the microbiome and the HIV reservoir. We analyzed HIV-1 DNA and RNA levels in isolated circulating CD4^+^ T cells, which reflect the size of the HIV reservoir ^44^. We found that *Firmicutes bacterium CAG 95* negatively correlated with CA-HIV-DNA levels and that the relation between *R. lactatiformans* and CA-HIV-RNA level was positive, while *Prevotella* species showed negative associations with CA-HIV-DNA and CA-HIV-RNA. How these bacterial species modulate the HIV reservoir remains speculative. *R. lactatiformans* has been linked with increased colonic IFN-γ^+^ T cells and immune activation ^45^, while a decreased abundance of *Firmicutes bacterium CAG 95* was found in subjects with hepatic steatosis ^46^. A previous study identified several immunogenic HLA–DR-presented *P. copri* peptides with the ability to induce T cells to produce the anti-viral cytokine IFN-γ ^47,48^. In our study, the *P. copri* healthy control-related strain was associated with decreased levels of IL-6 and TNF production, both of which can facilitate HIV-1 replication^49^.

We also investigated the role of HIV-associated gut dysbiosis in cytokine production capacity. HIV viral proteins can induce production of IL-10 by immune cells ^5^, which can be significantly reduced upon effective ART ^50^. A previous study isolated PBMCs from six untreated chronic infected PLHIV and six HCs and then cultured the PBMCs with bacterial lysates from type strains, with the PLHIV showing elevated levels of IL-10 in response to *P. copri* in comparison with HCs ^10^. In contrast, the PLHIV in our study received long-term ART, possibly resulting in a heterogeneous *P. copri* population that was negatively associated with IL-10 production capacity in the host. In addition to IL-10, HIV infection also induces priming of the monocyte IL-1β pathway in long-term treated PLHIV ^1^. Furthermore, our observations suggest that *B. vulgatus* was differentially associated with Pam3Cys-induced IL-1β production between PLHIV and HCs.

*P. copri* is not a monospecific taxonomic group and has many distinct clades with different immune functions ^51^. We therefore further investigated whether distinct *P. copri* strains were associated with different cytokine production and identified two *P. copri* strains: an HIV-related strain that showed enrichment in PLHIV and a control-related strain that was enriched in HCs. Only the control-related strain showed a negative association with IL-10, IL-6 and TNF production capacity, with no association found with the HIV-related strain. However, even when virologically suppressed, long-term treated PLHIV have significantly higher production levels of IL-10, IL-6 and TNF compared to HCs ^1^, which contributes to the chronic inflammation, HIV replication and disease progression in PLHIV ^49,50,52^, suggesting that the control-related strain plays a potentially protective role. The control-related strain consistently showed a positive association with CD4 counts and CD4-recovery-abs, which also supports this strain’s beneficial role in chronic inflammation. A large proportion of PLHIV seem to have lost this control-related *P. copri* strain, possibly leading to higher levels of IL-10, IL-6 and TNF production. Furthermore, the negative association between the control-related strain and cytokine production was observed in PLHIV, suggesting the altered immune system of long-term treated PLHIV may contribute to the strain–cytokine association. Mechanistically, five kinds of *P. copri* peptides have been found to induce IFN-γ production by T cells ^47^, with one peptide showing enrichment in the control-related strain that may contribute to its immune function.

A limitation of this study is potential batch effects among the different cohorts due to non-biological factors such as technical differences. We also did not have an independent cohort of PLHIV to validate our findings, but we could replicate the DI and FI scores in the 500FG cohort (built in the same medical center as the PLHIV cohort). Additionally, data on sexual behavior was only available for PLHIV and not for HCs. Most of the PLHIV were MSM, so our conclusions may not be generalizable to all PLHIV. To accurately indicate the effects on the gut microbiome of HIV itself vs. sexual practice, it is best to select HCs that consist of mainly MSM. Finally, no IL-10 production data were available from HCs, making it hard to compare the changes in IL-10 production capacity between PLHIV and HCs.

In conclusion, we observed differential microbial composition and function on species- and strain-level in long-term treated PLHIV. This HIV-associated bacterial signature was linked with HIV reservoir parameters and with PBMC production capacity of IL-1β and IL-10. A large fraction of the PLHIV have lost the control-related *P. copri* strain that was associated with IL-10, IL-6 and TNF production capacity and with CD4 counts and CD4-recovery-abs. The loss of this control-related strain may contribute to a higher level of IL-10, IL-6 and TNF production in PLHIV and to later immune activation and dysfunction. Our observation has provided deeper insight into the critical role of the gut microbiome during HIV infection. In the near future, the *P. copri* strain may be used as part of treatment for chronic inflammation, particularly cytokine imbalance.

## Materials and methods

### Study cohorts

The HIV cohort used in this study was described in our previous study ^1^ in which we recruited 211 PLHIV from the HIV clinic of the Radboud University Medical Center between December 2015 and February 2017. All participants were Caucasian individuals from the Netherlands. Study participants self-collected stool at their homes no more than 24h prior to study visits and stored the specimens in a refrigerator until being brought in for their visits. For this study, 143 metagenomic sequencing samples were available. Participants were excluded if they reported any antibiotics usage in the 3 months prior to fecal sample collection. One fecal sample was analyzed per individual. We included 190 age- and sex-matched HCs from the DMP cohort as the control group^53^. To replicate our findings in independent cohort, we also included 173 sex-matched HCs form 500FG cohort ^22^.

### Metagenomic data generation and profiling

The same protocol for fecal DNA isolation and metagenomic sequencing was used for both HIV samples and heathy control samples. Fecal DNA isolation was performed using the QIAamp Fast DNA Stool Mini Kit (Qiagen; cat. 51604). Fecal DNA was sent to Novogene to conduct library preparation and perform whole-genome shotgun sequencing on the Illumina HiSeq platform. Low quality reads and reads belonging to human genome were removed by mapping the data to the human reference genome (version HCBI37) using KneadData (v0.7.4). After filtering, the average read depth was 26.8 million for 143 HIV samples, 23.1 million for 190 DMP samples and 25.1 million for 173 500FG samples. Microbial taxonomic and functional profiles were determined using Metaphlan3 (v3.0.7) ^38^ and HUMAnN3 (v3.0.0.alpha.3) ^38^. The reads identified by MetaPhlAn3 are mapped to species-specific pangenomes with UniRef90 annotations, and the MetaPhlAn3-unclassified reads are translated and aligned to a protein database. Bacteria/pathways present in < 20% of the samples from one cohort were discarded.

### Strain profiles and analysis

We used Pangenome-based Phylogenomic Analysis3 (PanPhlAn3) ^38^ to identify the gene composition at the strain level. A total of 4,971 gene families from the *P. copri* pangenome were detected across 254 samples from the three cohorts. After filtering out the gene families that appeared in < 20% of all samples, 4260 gene families were studied in the subsequent analysis. A Jaccard distance matrix was built according to the presence/absence pattern of gene families. The strain cluster tree was constructed using the R basic function *hclust* with the hierarchical clustering method “complete”. Three strain clusters were defined at a tree height of 0.4. Visualizations were generated using the *dendextend* R package. Differentially abundant gene families were obtained using a logistic regression model, controlling for sex, age and read counts. The subsequent GO enrichment analysis was conducted using the *clusterProfiler* R package (v. 3.18.1) (*P. copri* GO annotation from PhanPhlan3), where the p.value for enrichment can be calculated by hypergeometric distribution. In the association analysis between cytokine production and HIV-related parameters and the two *P. copri* strains, we used linear regression, controlling for age, sex, read counts and sexual behavior, and using FDR < 0.1 as the significant threshold. We identified five *P. copri* peptides with immune function in IEDB ^54^ (Supplementary Table 23) and checked their abundance across the two strains in PLHIV and HCs using ShortBRED ^55^ using linear regression and controlling for age, sex and read counts.

### Microbial compositional and differential abundance analysis

The relative abundance data obtained from MetaPhlan3 was used to calculate bacterial diversity using the vegan R package (v. 2.5-7). Alpha diversity was calculated using the *diversity* function. The Bray-Curtis distance n-by-n matrix was built using the *vegdist* function, and then the PERMANOVA statistical test was applied on the matrix. We used the PCoA method to visualize the dissimilarities of beta diversity between different cohorts. To obtain the differentially abundant bacterial species and pathways between PLHIV and HCs, we first transformed the relative abundance data using CLR-transformation, as described before ^53^. Bacteria/pathways present in < 20% samples in at least one cohort were then discarded, and the remaining data were inverse-rank-transformed to follow a normal distribution. A linear regression model was then fitted, controlling for BMI, smoking status and read counts. Benjamini-Hochberg correction was used to correct for multiple hypothesis testing (using FDR < 0.05 as the significance threshold). DI and FI scores were calculated as the log2 ratio between geometric means of relative abundances of species/pathways that were enriched in PLHIV (linear regression FDR < 0.05 and HIV cohort beta > 0 in PLHIV vs. HCs) and depleted in PLHIV (linear regression FDR < 0.05 and HIV cohort beta <0 in PLHIV vs. HCs). DI and FI scores for HCs from the 500FG cohort were calculated using the same species/pathways from the comparison between the HIV and DMP cohorts.

### Measurement and analysis of *ex vivo* PBMC cytokine production

The detailed methods of *ex vivo* PBMC stimulation and cytokines measurements have been described before.^1^ In short, density centrifugation was performed on freshly collected venous blood to obtain the isolation of PBMCs. The freshly isolated cells were then incubated with different bacterial, fungal and viral stimuli at 37°C and 5% CO_2_ for either 24 hours or 7 days. IL-1β, IL-6, IL-1Ra, IL-10 and TNF were determined in the supernatants of the 24-hour PBMC or monocyte stimulation experiments using ELISAs. IL-17, IL-22 and IFN-γ were measured after the 7-day stimulation of PBMCs. Cytokine production data for the 500FG cohort was obtained using the same method, and the measurements that overlapped with those in the HIV cohort are summarized in the Supplementary Table 9. For the comparison of cytokine production between PLHIV and HCs from 500FG, we used different samples compared with the previous studies ^1,36^, so we conducted a re-analysis. A linear regression model was used, with correction for sex and age, using FDR < 0.05 as the significant threshold. Differentially abundant cytokine production and eight kinds of anti-inflammatory cytokine production (IL-10 and IL-1Ra) were included in the subsequent analysis.

### Microbial associations to HIV-related phenotypes and cytokine production capacity

For association analysis between gut microbiome and HIV-related phenotypes and cytokine production capacity, we included bacterial alpha diversity (Shannon index), beta diversity, P/B ratio, DI and FI score, as well as 76 species and 163 pathways that were significantly different between PLHIV and HCs from DMP cohort. Before Spearman correlation analysis, we first inverse-rank-transformed the data to follow a standard normal distribution, then adjusted all phenotypes for confounding factors (age, sex and read counts), and additionally adjusted for SO and Num-P, with FDR < 0.1 as the significant threshold.

## Supplemental information

Supplemental materials are available.

## Acknowledgements

We thank all the volunteers in the 200HIV cohort, DMP cohort and 500FG cohort for their participation and the project staff for their help and management. This research was funded by Dutch Heart Foundation IN-CONTROL (CVON2018-27, to A.Z., L.J., M.G.N., and J.F.). J.F. is also supported by the ERC Consolidator grant (grant agreement No. 101001678), NWO-VICI grant VI.C.202.022, and the Netherlands Organ-on-Chip Initiative, an NWO Gravitation project (024.003.001) funded by the Ministry of Education, Culture and Science of the government of The Netherlands. A.Z. is supported the ERC Starting Grant 715772, NWO-VIDI grant 016.178.056, and the NWO Gravitation grant Exposome-NL (024.004.017). M.G.N was supported by an ERC Advanced Grant and a Spinoza Grant of the Netherlands Organization for Scientific Research. R.K.W. is supported by the Seerave Foundation and the Dutch Digestive Foundation (16-14). Y.Z. is supported by a joint fellowship from the University Medical Centre Groningen and China Scholarship Council (CSC202006170040). We also thank the Genomics Coordination Center for providing data infrastructure and access to high performance computing clusters, Kate Mclntyre for critical reading and editing and Maartje Cleophas for data management and transfer.

## Author contributions

J.F. and A.v.d.V. conceptualized and managed the study. W.v.d.H., L.V., Q.d.M., L.A.B.J., R.K.W., A.Z. and M.G.N. contributed to data generation. Y.Z, S.A., D.W. and R.G. analyzed the data. Y.Z., J.F. and A.v.d.V. drafted the manuscript. S.A., N.V., D.W., V.M., W.v.d.H., R.G., R.K.W., A.Z., L.V., Q.d.M., L.A.B.J., and M.G.N. reviewed and edited the manuscript.

## Correspondence contact

Correspondence should be addressed to

J.F. and A.v.d.V.

## Ethical approval

The 200HIV study was approved by the Medical Ethical Review Committee region Arnhem-Nijmegen (CMO2012-550). For the Dutch Microbiome Project (DMP) cohort, the Lifelines study was approved by the medical ethical committee from the University Medical Center Groningen (METc number: 2017/152). The 500 Functional Genomics (500FG) study was approved by the Ethical Committee of Radboud University Nijmegen (NL42561.091.12, 2012/550). All informed consents were collected for all participants. Experiments were conducted in accordance with the principles of the Declaration of Helsinki.

## Data Availability

All relevant data supporting the key findings of this study are available within the article and its Supplementary Information files. The raw metagenomic sequencing data of all subjects were all publicly available: the 200HIV cohort via NCBI Short Read Archive (SRA) under accession number PRJNA820547 (BioProject), the DMP cohort via the European Genome-Phenome Archive under accession number EGAS00001005027 and the 500FG cohort via SRA under accession number PRJNA319574 (BioProject). Due to informed consent regulation, the clinical data of the 200HIV cohort, the 500FG cohort and the DMP cohort are available upon request to Radboud University Medical Center and the LifeLines, respectively. This includes the submission of a letter of intention to the corresponding data access committee: the data access committee for the 200HIV cohort (Maartje Jacob-Cleophas), the Lifelines Data Access Committee for the Dutch Microbiome Project (https://forms.gle/eHeBdXJMXbVvCJRc8), and the Human Functional Genomics Data Access Committee for 500FG (Martin Jaeger). Data sets can be made available under a data transfer agreement and the data usage access is subject to local rules and regulations.

## Code availability

For this study, the following software was used: KneadData (v0.7.4), Bowtie2 (v2.3.4.2), MetaPhlAn2 (v3.0.7), HUMAnN2 (v 3.0.0.alpha.3), PanPhlAn (v 3.1) and shortBRED (v0.9.5). Code used for the statistical analyses is publicly available at GitHub: https://github.com/White-Shinobi/HIV-and-gut-microbiome.

## References

1. van der Heijden, W. A. et al. Chronic HIV infection induces transcriptional and functional reprogramming of innate immune cells. JCI insight 6, (2021).

2. Hileman, C. O. & Funderburg, N. T. Inflammation, Immune Activation, and Antiretroviral Therapy in HIV. Curr. HIV/AIDS Rep. 14, 93–100 (2017).

3. Keating, S. M., Jacobs, E. S. & Norris, P. J. Soluble mediators of inflammation in HIV and their implications for therapeutics and vaccine development. Cytokine Growth Factor Rev. 23, 193–206 (2012).

4. Osuji, F. N., Onyenekwe, C. C., Ahaneku, J. E. & Ukibe, N. R. The effects of highly active antiretroviral therapy on the serum levels of pro-inflammatory and anti-inflammatory cytokines in HIV infected subjects. J. Biomed. Sci. 25, 88 (2018).

5. Planès, R., Serrero, M., Leghmari, K., BenMohamed, L. & Bahraoui, E. HIV-1 Envelope Glycoproteins Induce the Production of TNF-α and IL-10 in Human Monocytes by Activating Calcium Pathway. Sci. Rep. 8, 17215 (2018).

6. Klatt, N. R., Chomont, N., Douek, D. C. & Deeks, S. G. Immune activation and HIV persistence: implications for curative approaches to HIV infection. Immunol. Rev. 254, 326–342 (2013).

7. Crakes, K. R. & Jiang, G. Gut Microbiome Alterations During HIV/SIV Infection: Implications for HIV Cure. Front. Microbiol. 10, 1104 (2019).

8. Vujkovic-Cvijin, I. & Somsouk, M. HIV and the Gut Microbiota: Composition, Consequences, and Avenues for Amelioration. Curr. HIV/AIDS Rep. 16, 204–213 (2019).

9. Vázquez-Castellanos, J. F. et al. Interplay between gut microbiota metabolism and inflammation in HIV infection. ISME J. 12, 1964–1976 (2018).

10. Lozupone, C. A. et al. Alterations in the gut microbiota associated with HIV-1 infection. Cell Host Microbe 14, 329–339 (2013).

11. Ray, S. et al. Altered Gut Microbiome under Antiretroviral Therapy: Impact of Efavirenz and Zidovudine. ACS Infect. Dis. 7, 1104–1115 (2021).

12. Pinto-Cardoso, S. et al. Fecal Bacterial Communities in treated HIV infected individuals on two antiretroviral regimens. Sci. Rep. 7, 43741 (2017).

13. Tuddenham, S., Koay, W. L. & Sears, C. HIV, Sexual Orientation, and Gut Microbiome Interactions. Dig. Dis. Sci. 65, 800–817 (2020).

14. Armstrong, A. J. S. et al. An exploration of Prevotella-rich microbiomes in HIV and men who have sex with men. Microbiome 6, 198 (2018).

15. Vujkovic-Cvijin, I. et al. Dysbiosis of the gut microbiota is associated with HIV disease progression and tryptophan catabolism. Sci. Transl. Med. 5, 193ra91 (2013).

16. Vujkovic-Cvijin, I. et al. HIV-associated gut dysbiosis is independent of sexual practice and correlates with noncommunicable diseases. Nat. Commun. 11, 2448 (2020).

17. Ishizaka, A. et al. Unique Gut Microbiome in HIV Patients on Antiretroviral Therapy (ART) Suggests Association with Chronic Inflammation. Microbiol. Spectr. 9, e0070821 (2021).

18. El-Far, M. et al. Upregulated IL-32 Expression And Reduced Gut Short Chain Fatty Acid Caproic Acid in People Living With HIV With Subclinical Atherosclerosis. Front. Immunol. 12, 664371 (2021).

19. Dillon, S. M. et al. HIV-1 infection of human intestinal lamina propria CD4+ T cells in vitro is enhanced by exposure to commensal Escherichia coli. J. Immunol. 189, 885–896 (2012).

20. Dillon, S. M. et al. Enhancement of HIV-1 infection and intestinal CD4+ T cell depletion ex vivo by gut microbes altered during chronic HIV-1 infection. Retrovirology 13, 5 (2016).

21. Dillon, S. M. et al. Low abundance of colonic butyrate-producing bacteria in HIV infection is associated with microbial translocation and immune activation. AIDS 31, 511–521 (2017).

22. Schirmer, M. et al. Linking the Human Gut Microbiome to Inflammatory Cytokine Production Capacity. Cell 167, 1897 (2016).

23. Gootenberg, D. B., Paer, J. M., Luevano, J.-M. & Kwon, D. S. HIV-associated changes in the enteric microbial community: potential role in loss of homeostasis and development of systemic inflammation. Curr. Opin. Infect. Dis. 30, 31–43 (2017).

24. Nowak, P. et al. Gut microbiota diversity predicts immune status in HIV-1 infection. AIDS 29, 2409–2418 (2015).

25. Zilberman-Schapira, G. et al. The gut microbiome in human immunodeficiency virus infection. BMC Med. 14, 83 (2016).

26. Vázquez-Castellanos, J. F. et al. Altered metabolism of gut microbiota contributes to chronic immune activation in HIV-infected individuals. Mucosal Immunol. 8, 760–772 (2015).

27. Gosmann, C. et al. Lactobacillus-Deficient Cervicovaginal Bacterial Communities Are Associated with Increased HIV Acquisition in Young South African Women. Immunity 46, 29–37 (2017).

28. Daillère, R. et al. Enterococcus hirae and Barnesiella intestinihominis Facilitate Cyclophosphamide-Induced Therapeutic Immunomodulatory Effects. Immunity 45, 931–943 (2016).

29. Murray, M. F. Tryptophan depletion and HIV infection: a metabolic link to pathogenesis. Lancet. Infect. Dis. 3, 644–652 (2003).

30. Papadia, C. et al. Plasma citrulline as a quantitative biomarker of HIV-associated villous atrophy in a tropical enteropathy population. Clin. Nutr. 29, 795–800 (2010).

31. Baumgartner, M. R. et al. Hyperammonemia with reduced ornithine, citrulline, arginine and proline: a new inborn error caused by a mutation in the gene encoding delta(1)-pyrroline-5-carboxylate synthase. Hum. Mol. Genet. 9, 2853–2858 (2000).

32. Dirajlal-Fargo, S. et al. Comprehensive assessment of the arginine pathway and its relationship to inflammation in HIV. AIDS 31, 533–537 (2017).

33. Smith, E. & Morowitz, H. J. Universality in intermediary metabolism. Proc. Natl. Acad. Sci. U. S. A. 101, 13168–13173 (2004).

34. Ling, Z. et al. Alterations in the Fecal Microbiota of Patients with HIV-1 Infection: An Observational Study in A Chinese Population. Sci. Rep. 6, 30673 (2016).

35. Dillon, S. M. et al. An altered intestinal mucosal microbiome in HIV-1 infection is associated with mucosal and systemic immune activation and endotoxemia. Mucosal Immunol. 7, 983–994 (2014).

36. Van de Wijer, L. et al. The Architecture of Circulating Immune Cells Is Dysregulated in People Living With HIV on Long Term Antiretroviral Treatment and Relates With Markers of the HIV-1 Reservoir, Cytomegalovirus, and Microbial Translocation. Front. Immunol. 12, 661990 (2021).

37. Scholz, M. et al. Strain-level microbial epidemiology and population genomics from shotgun metagenomics. Nat. Methods 13, 435–438 (2016).

38. Beghini, F. et al. Integrating taxonomic, functional, and strain-level profiling of diverse microbial communities with biobakery 3. Elife 10, (2021).

39. Chang, Y.-J. et al. Complete genome sequence of Acidaminococcus fermentans type strain (VR4). Stand. Genomic Sci. 3, 1–14 (2010).

40. Parada Venegas, D. et al. Short Chain Fatty Acids (SCFAs)-Mediated Gut Epithelial and Immune Regulation and Its Relevance for Inflammatory Bowel Diseases. Front. Immunol. 10, 277 (2019).

41. Schröder, C., Matthies, A., Engst, W., Blaut, M. & Braune, A. Identification and expression of genes involved in the conversion of daidzein and genistein by the equol-forming bacterium Slackia isoflavoniconvertens. Appl. Environ. Microbiol. 79, 3494–3502 (2013).

42. Zagato, E. et al. Endogenous murine microbiota member Faecalibaculum rodentium and its human homologue protect from intestinal tumour growth. Nat. Microbiol. 5, 511–524 (2020).

43. Churchill, M. J., Deeks, S. G., Margolis, D. M., Siliciano, R. F. & Swanstrom, R. HIV reservoirs: what, where and how to target them. Nature reviews. Microbiology vol. 14 55–60 (2016).

44. Rutsaert, S., De Spiegelaere, W., De Clercq, L. & Vandekerckhove, L. Evaluation of HIV-1 reservoir levels as possible markers for virological failure during boosted darunavir monotherapy. J. Antimicrob. Chemother. 74, 3030–3034 (2019).

45. Frankel, A. E. et al. Cancer Immune Checkpoint Inhibitor Therapy and the Gut Microbiota. Integr. Cancer Ther. 18, 1534735419846379 (2019).

46. Zeybel, M. et al. Multi-omics analysis reveals the impact of microbiota on host metabolism in hepatic steatosis. medRxiv +x2021.05.22.21257482 (2021) doi:10.1101/2021.05.22.21257482.

47. Pianta, A. et al. Identification of Novel, Immunogenic HLA-DR-Presented Prevotella copri Peptides in Patients With Rheumatoid Arthritis. Arthritis Rheumatol. (Hoboken, N.J.) 73, 2200–2205 (2021).

48. Kak, G., Raza, M. & Tiwari, B. K. Interferon-gamma (IFN-γ): Exploring its implications in infectious diseases. Biomol. Concepts 9, 64–79 (2018).

49. Connolly, N. C., Riddler, S. A. & Rinaldo, C. R. Proinflammatory cytokines in HIV disease-a review and rationale for new therapeutic approaches. AIDS Rev. 7, 168–180 (2005).

50. Brockman, M. A. et al. IL-10 is up-regulated in multiple cell types during viremic HIV infection and reversibly inhibits virus-specific T cells. Blood 114, 346–356 (2009).

51. Iljazovic, A., Amend, L., Galvez, E. J. C., de Oliveira, R. & Strowig, T. Modulation of inflammatory responses by gastrointestinal Prevotella spp. - From associations to functional studies. Int. J. Med. Microbiol. 311, 151472 (2021).

52. Mahajan, S. D. et al. Role of chemokine and cytokine polymorphisms in the progression of HIV-1 disease. Biochem. Biophys. Res. Commun. 396, 348–352 (2010).

53. Gacesa, R. et al. The Dutch Microbiome Project defines factors that shape the healthy gut microbiome. bioRxiv 1–33 (2020).

54. Vita, R. et al. The Immune Epitope Database (IEDB): 2018 update. Nucleic Acids Res. 47, D339–D343 (2019).

55. Kaminski, J. et al. High-Specificity Targeted Functional Profiling in Microbial Communities with ShortBRED. PLoS Comput. Biol. 11, e1004557 (2015).

