## Supplemental Figures for "HIV-linked gut dysbiosis associates with cytokine production capacity in viral-suppressed people living with HIV"

**Figure S1. Comparison of the microbial pathway alpha diversity between people living with HIV (PLHIV) and healthy controls (HCs) from DMP cohort.** Related to Figure 1. Y-axis is the Shannon index at the bacterial pathway level.

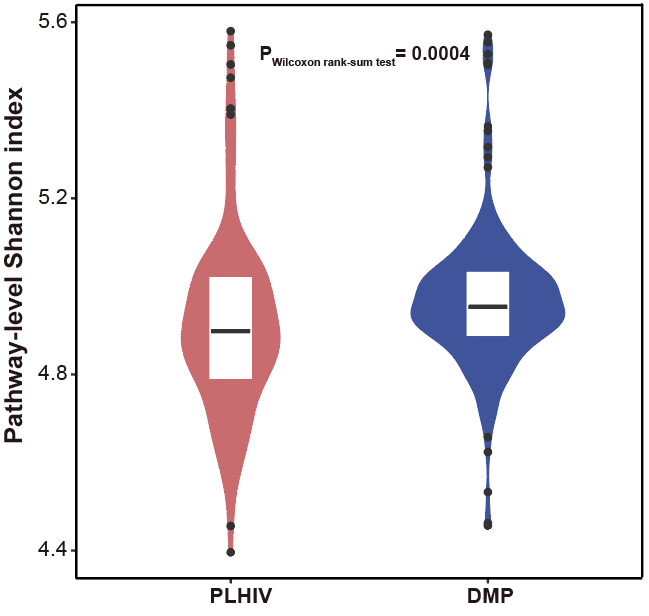

**Figure S2. Relative abundance of *Prevotella* species in PLHIV and HCs in DMP cohort.** Related to Figure 1. **a.** *Prevotella copri*. **b.** *Prevotella sp 885.* **c.** *Prevotella sp AM42 24*. **d.** *Prevotella sp CAG 1092*. **e.** *Prevotella sp CAG 279*. **f.** *Prevotella sp CAG 520*. **g.** *Prevotella sp CAG 5226*. **h.** *Prevotella stercorea.*

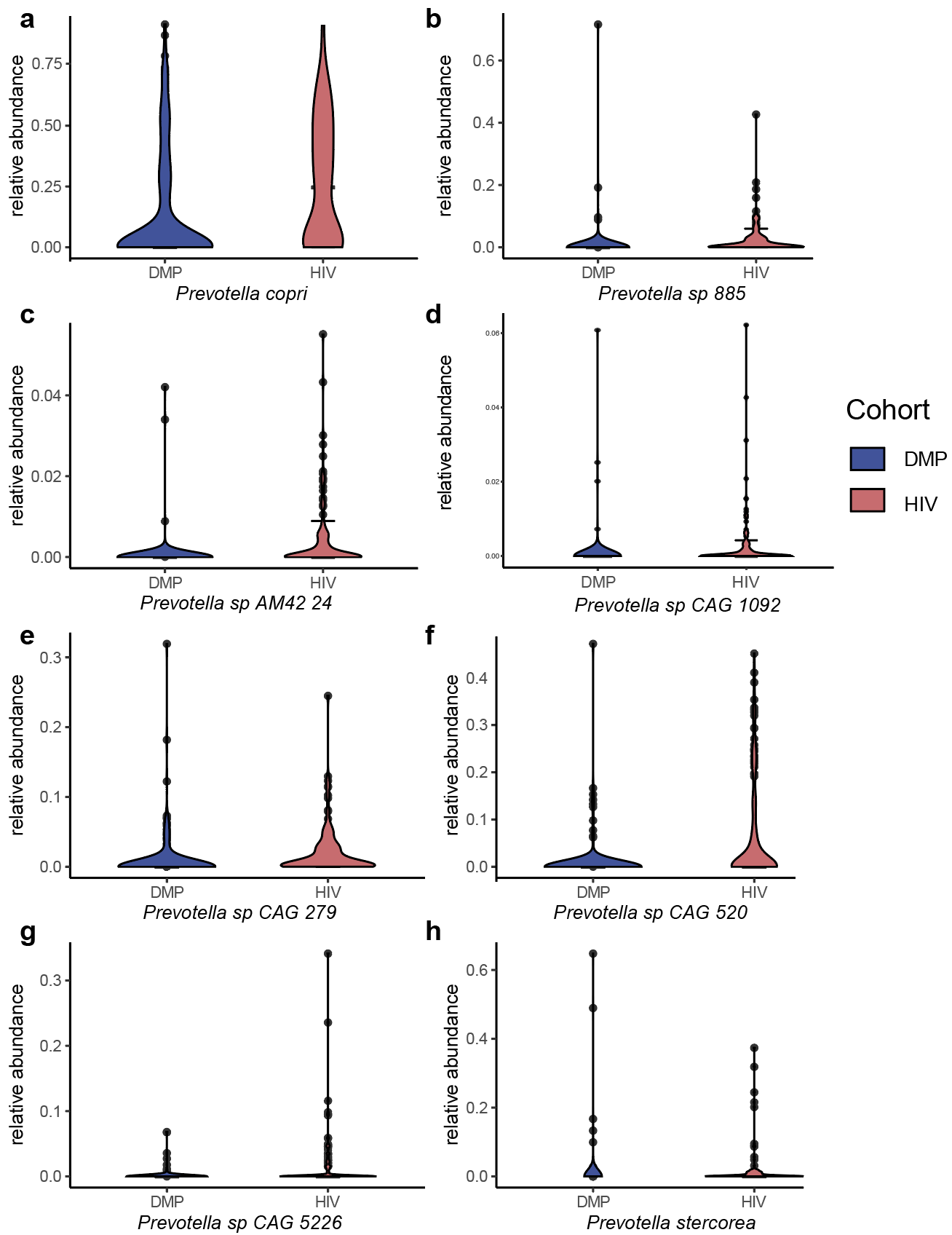

**Figure S3. Violin plots comparing microbiome indices between the 500FG, DMP and HIV cohorts. a.** Ratio between *Prevotella* and *Bacteroides* genus (P/B). **b.** Dysbiosis index (DI) score. **c.** Functional index (FI) score.

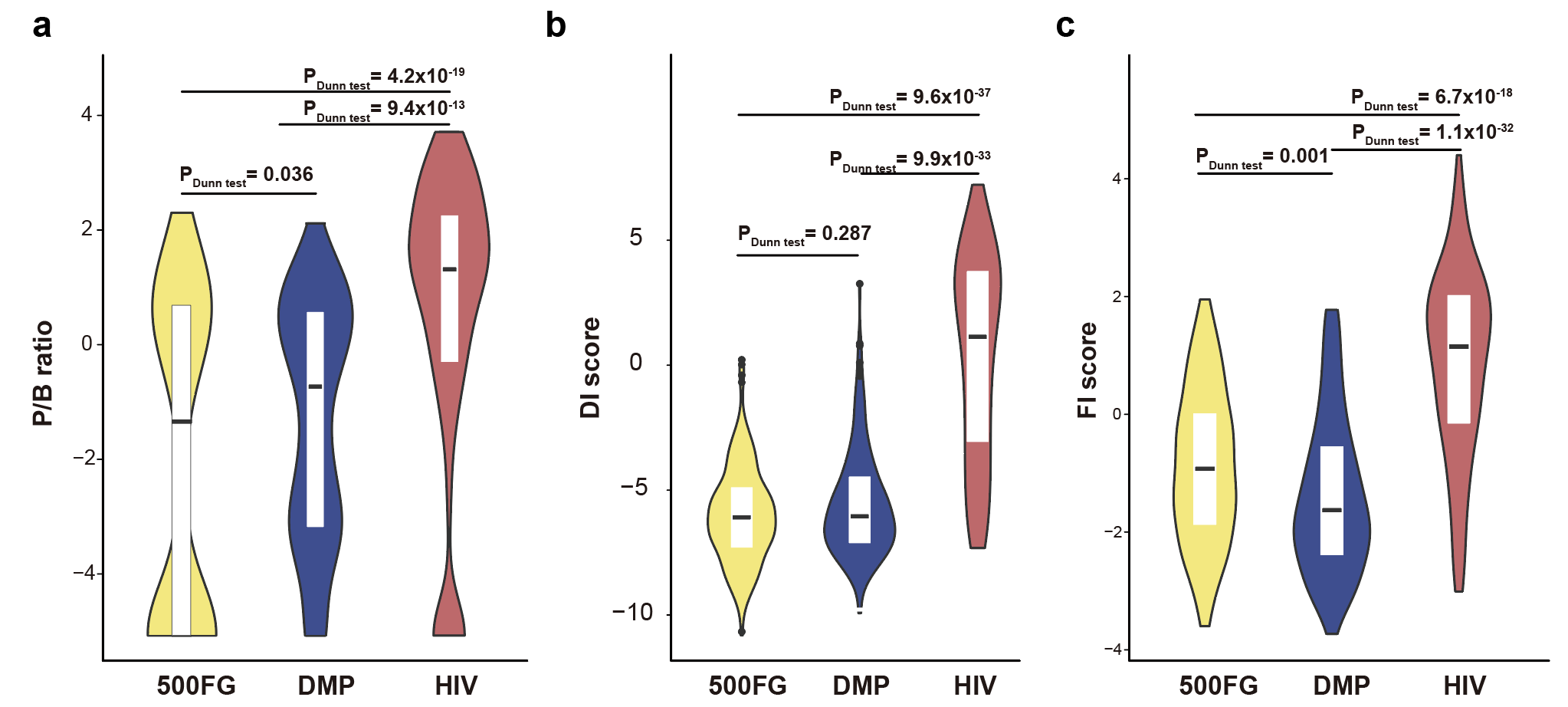

**Figure S4. Distribution of HIV-related phenotypes in PLHIV.**

**a-c.** HIV-1 reservoir measurements in circulating CD4^+^ T cells. **(a)** CD4^+^ T cell–associated HIV-1 RNA (CA-HIV-RNA), **(b)** CD4^+^ T cell–associated HIV-1 DNA (CA-HIV-DNA), **(c)** ratio between CA-RNA and CA-DNA (CA-RNA/CA-DNA). **d-h.** CD4^+^ T cell counts. **(d)** CD4^+^ T cell counts nadir (CD4 nadir), **(e)** CD4^+^ T cell counts latest (CD4 counts), **(f)** ratio between CD4^+^ T cell count and CD8^+^ T cell counts (CD4/CD8), **(g)** CD4^+^ cell recovery absolute value (CD4-recovery-abs), **(h)** CD4^+^ cell recovery relative value (CD4-recovery-re). **i.** Sexual orientation data and receptive anal intercourse (RAI) data. **j-k.** Plasma viral load. **(j)** Zenith value of HIV RNA load level in plasma (HIV RNA zenith), **(k)** HIV RNA level in plasma (HIV RNA load). **l-n.** Duration related to infection and treatment. **(l)** Duration of HIV infection (HIV duration), **(m)** Duration of combined antiretroviral therapy (cART duration), **(n)** Duration between diagnosis of HIV and the start of cART (Time diagnosis-cART). Y-axis in A–H and J–N refers to the count of the phenotypes.

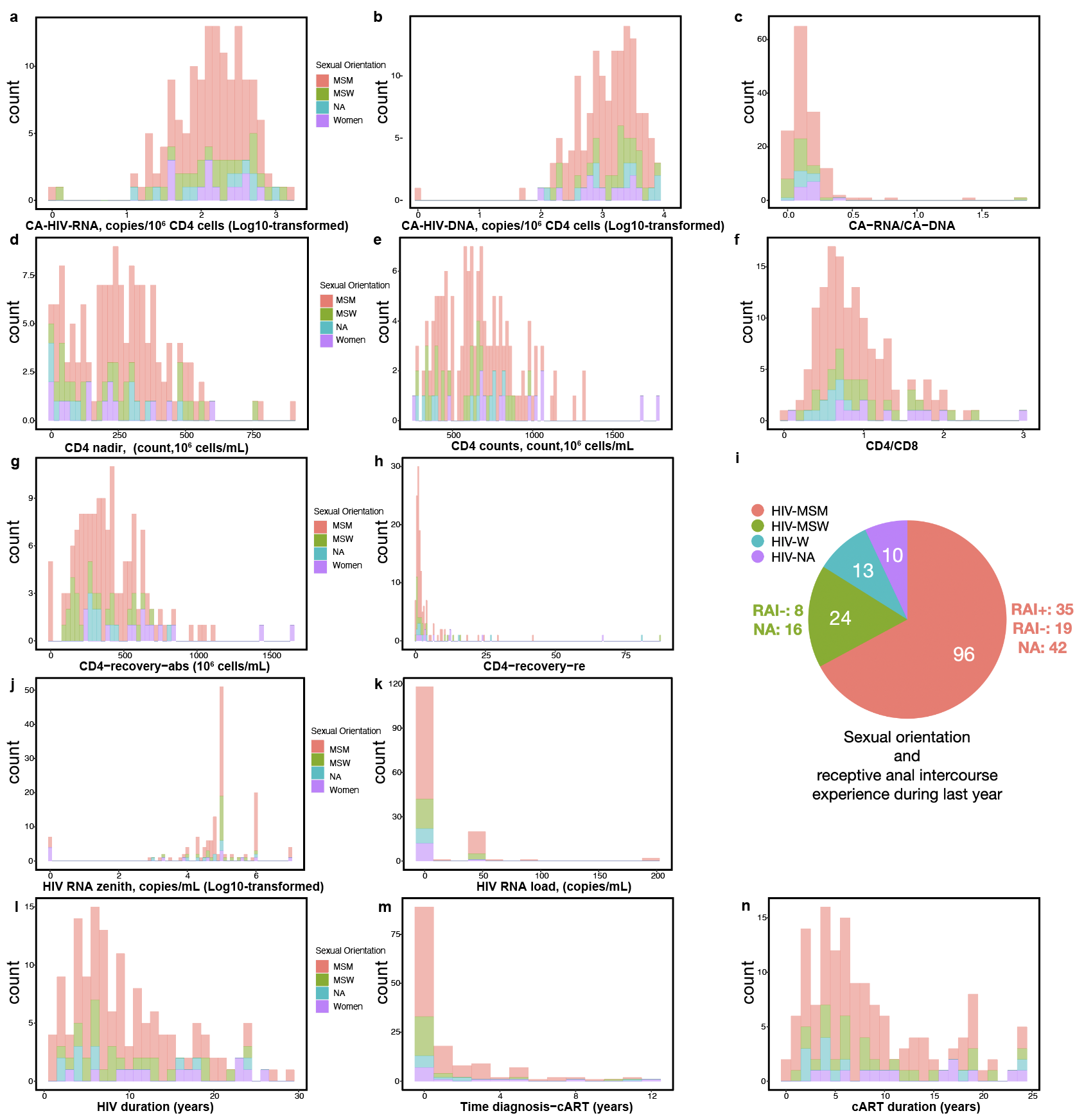

**Figure S5. Relationships between HIV-related phenotypes.** Heatmap shows the Spearman correlation rho between HIV-related phenotypes. Box color indicates the correlation rho. White boxes indicate an FDR value for the correlation > 0.1.

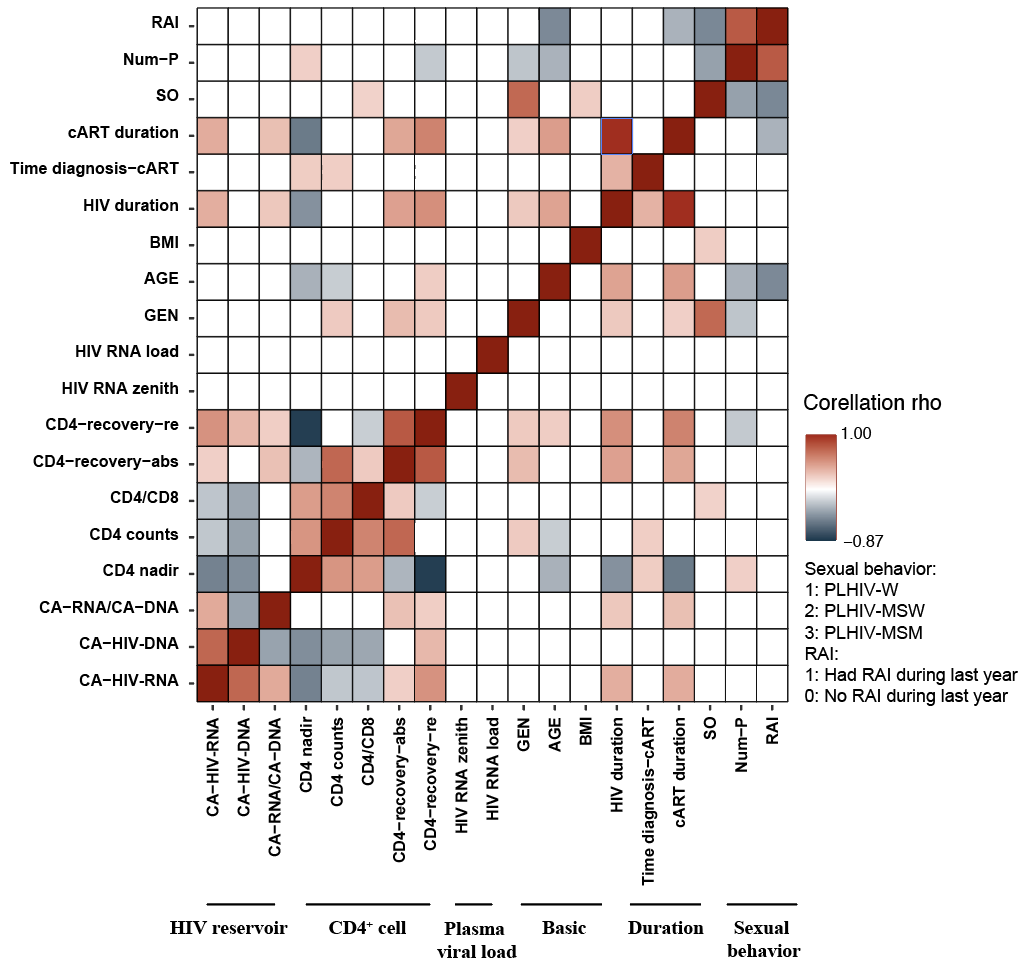

**Figure S6. Heatmap of associations between HIV-associated bacterial signatures (Shannon index, beta diversity, P/B ratio, DI and FI score) and HIV-related phenotypes.** Heatmap shows Spearman correlation rho. HIV-related phenotypes are corrected for age, gender, read count and sexual behavior. Box color indicates the Spearman correlation rho. White box indicates P > 0.05. Bray-Curtis distance of species and then multiplied by ten to rescale.

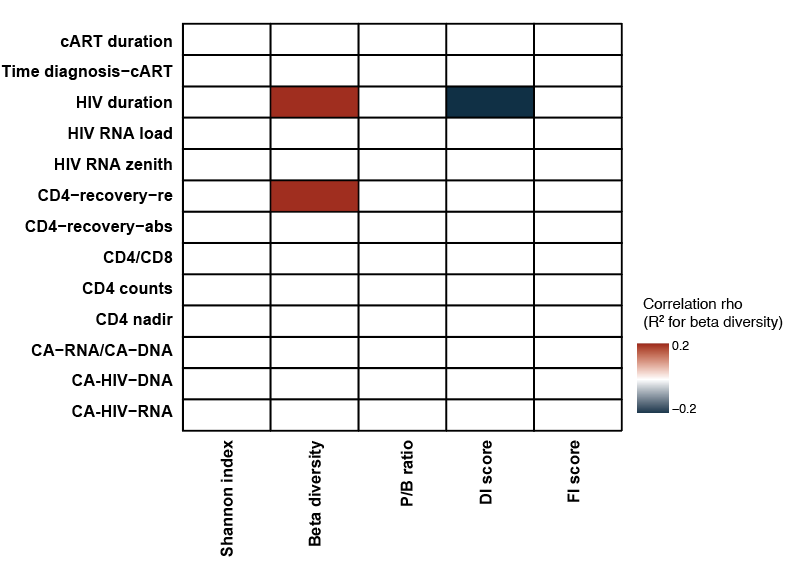

**Figure S7. Relationship between HIV-related phenotypes and cytokine production capacity.** Heatmap shows the Spearman correlation rho between HIV-related phenotypes and cytokine production capacity. Box colors indicate the correlation rho. White indicates an FDR value for the correlation was > 0.1.

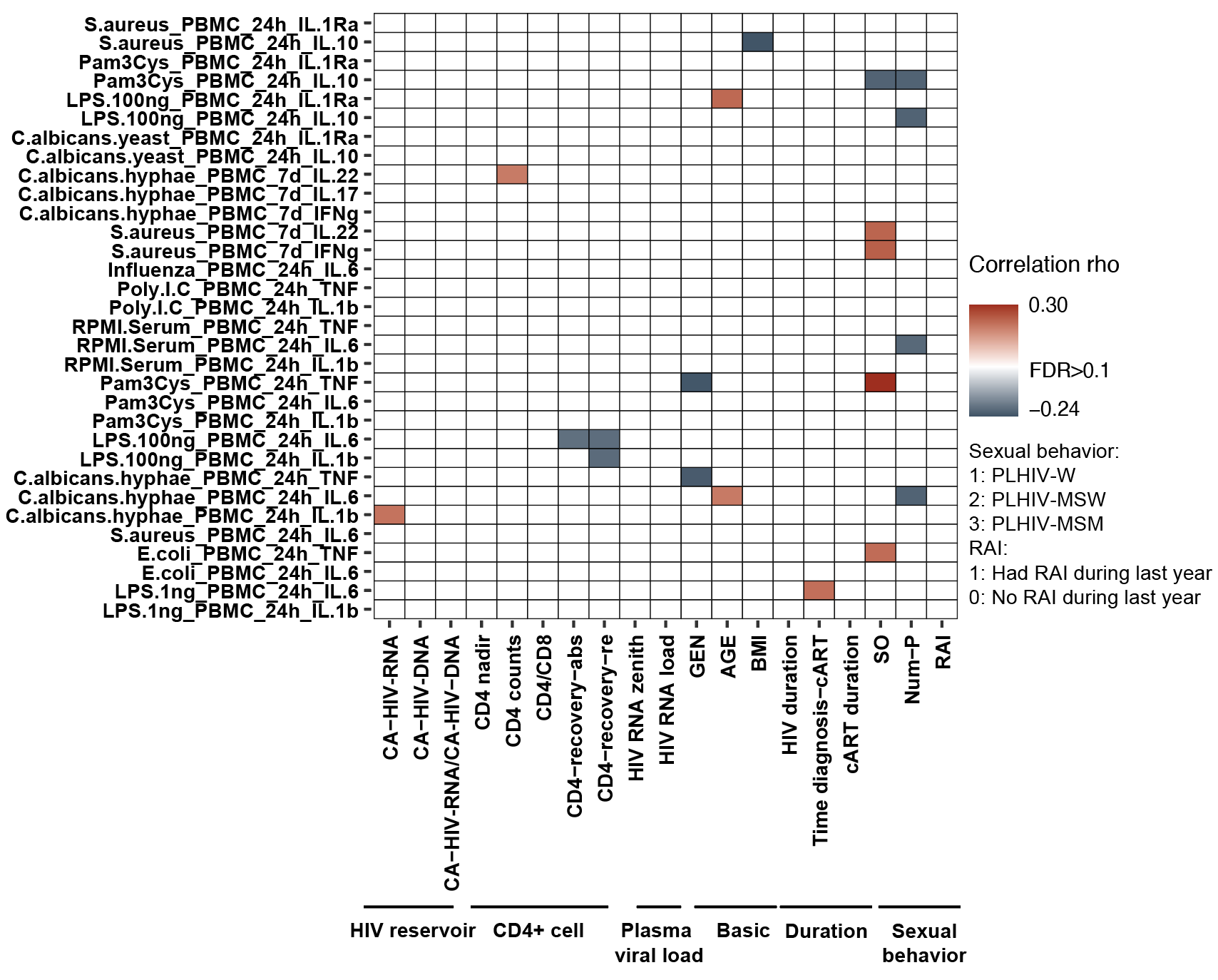

**Figure S8. Distinct associations of the relative abundances of the control-related strain and HIV-related *Prevotella copri* strain with IL-6 (a, b) and TNF production capacity (c) in HCs from 500FG cohort, established using linear regression.** Cytokine production was corrected for age, sex, and read counts.

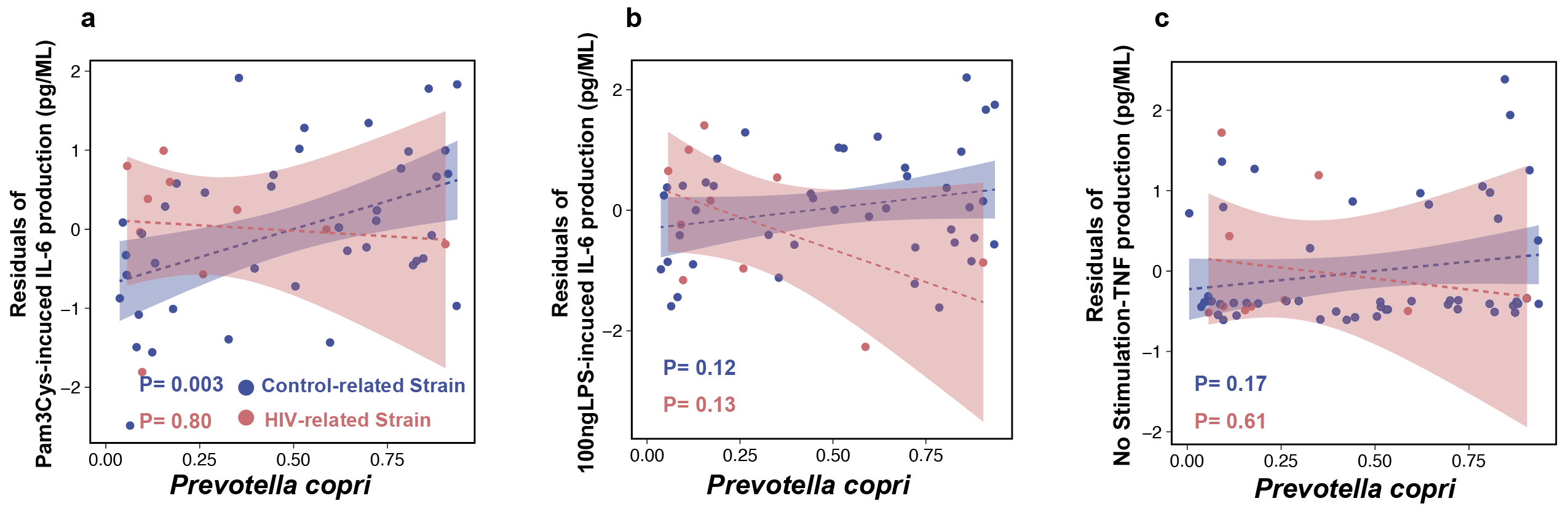
